# Development of the Left Arcuate Fasciculus is Linked to Learning Gains in Reading, but not Math

**DOI:** 10.1101/2024.11.07.622498

**Authors:** Ethan Roy, Emily M. Harriott, Tin Q. Nguyen, Adam Richie-Halford, Ariel Rokem, Laurie E. Cutting, Jason D. Yeatman

## Abstract

Past studies leveraging cross-sectional data have raised questions surrounding the relationship between diffusion properties of the white matter and academic skills. Some studies have suggested that white matter properties serve as static predictors of academic skills, whereas other studies have observed no such relationship. On the other hand, longitudinal studies have suggested that within-individual changes in the white matter are linked to learning gains over time. In the present study, we look to replicate and extend the previous longitudinal results linking longitudinal changes in the white matter properties of the left arcuate fasciculus to individual differences in reading development. To do so, we analyzed diffusion MRI data, along with reading and mathematics scores in a longitudinal sample of 340 students as they progressed from 1st grade into 4th grade. Longitudinal growth models revealed that year-to-year within-individual changes in reading scores, but not math, were related to the development of the left arcuate fasciculus. These findings provide further evidence linking the dynamics of white matter development and learning in a unique sample and highlight the importance of longitudinal designs.

## 1. Introduction

Over the course of the lifespan, an individual’s experiences shape the development of white matter tracts, the large bundles of axons connecting distinct cortical areas. Experience has been shown to sculpt the properties of these white matter tracts on the time scale of hours (Sagi et al., 2012) to weeks and months (Huber et al., 2018). The acquisition of academic skills that rely on culturally invented symbolic systems serves as a prominent example of experience-dependent white matter plasticity. Although the human brain has not evolved to innately process and manipulate symbolic representations of language and number, its incredible capacity plasticity allows it to adapt in response to repeated exposure to these cultural inventions to give rise to complex behaviors, such as reading and math (Caffarra et al., 2021; Dehaene, 2005; Dehaene & Cohen, 2007).

Past research leveraging diffusion MRI has led to mixed, and sometimes conflicting, conclusions about the relationship between white matter properties and academic skills. In the case of reading, some cross-sectional studies have suggested that higher fractional anisotropy (FA) in the left arcuate fasciculus, inferior longitudinal fasciculus (ILF), and superior longitudinal fasciculus (SLF) correlate with higher reading scores (Van Der Auwera et al., 2021; Vanderauwera et al., 2017; Welcome & Joanisse, 2014). However, other studies have observed the opposite relationship, with higher levels of reading skill corresponding to lower levels of FA these tracts (Yeatman et al., 2011) or no relationship whatsoever between reading skill and white matter properties (Meisler & Gabrieli, 2021; Moreau et al., 2018; Roy, Richie-Halford, et al., 2024).

These mixed findings suggest that past results reporting a cross-sectional relationship between academic skills and white matter may stem from idiosyncratic properties of the study sample. In fact, past studies incorporating measures of socioeconomic status into their analyses have demonstrated that the relationship between reading skill and white matter properties varies as a function of socioeconomic status (Gullick et al., 2016; Ozernov-Palchik et al., 2019; Turesky et al., 2022). Based on these findings, the reported link between reading skill and properties of the left arcuate and SLF may stem from characteristics of the sample populations, such as environmental, demographic or socioeconomic factors, and may not generalize to the broader population.

In the case of mathematics skills, FA in the left SLF, left ILF, and left arcuate has been shown to predict arithmetic ability (Jolles et al., 2016; Matejko et al., 2013; Matejko & Ansari, 2015; Tsang et al., 2009; Van Beek et al., 2014; van Eimeren et al., 2008). In addition to this set of left lateralized tracts, studies have also found that the diffusion properties of right hemisphere tracts also correlate with mathematics abilities (Cantlon et al., 2009; Polspoel et al., 2019; Rykhlevskaia et al., 2009; Till et al., 2011), suggesting that, unlike reading, the neural circuitry underlying mathematics skill includes a bilateral collection of white matter tracts. These findings again suggest that individual differences in white matter properties might relate to differences in mathematics performance. However, as with past studies of reading, these findings are based on small scale, cross-sectional, samples of convenience and have yet to be replicated using large, representative, multi-site consortium datasets.

In contrast to the mixed findings observed in cross-sectional studies, longitudinal and intervention studies using repeated measures to evaluate within-individual change over time have demonstrated a consistent relationship between changes in the white matter and the development of reading and math skill. Observational studies have shown that within-individual gains in reading skill are associated with the development of the left arcuate fasciculus (Myers et al., 2014; Roy, Richie-Halford, et al., 2024; Wang et al., 2017; Yeatman, Dougherty, Ben-Shachar, et al., 2012) and that delayed development of frontal-parietal and posterior parietal white matter networks is associated with difficulties in mathematics (Ranpura et al., 2013). Furthermore, targeted learning intervention studies have demonstrated that intensive educational experiences can drive changes in an individual’s white matter properties that are associated with behavioral gains in reading skills (Huber et al., 2018; Meisler et al., 2023) and in mathematics (Jolles et al., 2016; Klein et al., 2019). Together, these results suggest that white matter properties underlying reading skill change over time and, in fact, change in response to the educational opportunities and experiences afforded by an individual’s environment.

A recent study by Roy et al. (2024) aimed to clarify past findings and evaluate both the cross-sectional and longitudinal hypotheses surrounding the relationship between white matter and reading skill. In a sample of children aged 5-10 years, within-individual changes in the diffusion properties of the left arcuate related to within-individual gains in reading skill based on the PLING sample (Wierenga et al., 2018). Interestingly, these analyses did not find any cross-sectional differences between reading skill and white matter properties at a given time point and three other data sets totaling 14,249 participants confirmed the lack of a generalizable cross-sectional relationship between white matter and reading. Furthermore, models examining the temporal dynamics of reading gains and the development of the left arcuate suggested that improvements in reading skills precede changes in the white matter. Together, these results suggest that the white matter properties of the core reading circuitry change over time, potentially in response to environmental and educational factors that promote the development of reading skills.

In the present study, we look to replicate the findings reported in Roy et al. (2024) using unique diffusion MRI data collected from an independent longitudinal sample of children between six and eight years of age, from a completely different regional context in the United States (the Southern United States, whereas the PLING sample was collected in California). We use pyAFQ (Kruper et al., 2021) to examine the development of the left and right arcuate fasciculus and their relationship to the development of reading and mathematics skills. The use of pyAFQ to generate tract profiles allows us to leverage non-linear modeling approaches, such as generalized additive mixed-models (GAMM; Hastie & Tibshirani, 1987), to examine the development of white matter properties over the entire length of each tract, as opposed to the approach used in Roy et al. (2024), which relied on a single average diffusion metric for each tract. This approach to modeling tract profiles is appealing, as diffusion properties vary over the length of a tract and important group differences in these diffusion properties may be obscured by collapsing these data into a single value (Muncy et al., 2022; Yeatman et al., 2012).

Our analyses reveal that, within this longitudinal sample, within-individual changes in the mean diffusivity (MD) of the left arcuate fasciculus, but not the right arcuate, are related to within-individual changes in reading scores. There was no such relationship between development of the left arcuate and mathematics skill, suggesting that this relationship is specific to reading. To explain these findings we examined between-individual differences in the growth trajectories of both white matter and academic skills and found significant interindividual variation in both reading and white matter development, but not in math skill. Furthermore, node-wise modeling of the time series of MD development and reading development suggested that individual gains in reading skill precede changes in the white matter properties along the length of the left arcuate fasciculus. These findings serve to replicate past findings demonstrating a longitudinal relationship between reading skill and the development of the left arcuate and suggest that this longitudinal link is not specific to certain portions of the left arcuate, but rather extend across the length of the entire tract.

## 2. Methods

### 2.1 Participants

340 first-graders were recruited from the greater Nashville area to participate in this longitudinal study of the development of academic skills. Starting in the summer after first grade, participants completed annual behavioral and neuroimaging sessions in which they completed either the Woodcock-Johnson III or IV assessments of basic reading and mathematics (Woodcock et al., 2001), as well as structural and diffusion neuroimaging scans. Behavioral and neuroimaging data were collected across four waves. The first time point involved the collection of neuroimaging and/or behavioral data from 268 participants, at the second time point from 220 participants, at the third time point from 183 participants, and at the fourth time point from 92 participants.

At the onset of the study and at each observation, parents provided informed consent and participating children provided assent before data collection. Data collection was carried out with approval from the Vanderbilt University Institutional Review Board. Data sharing was carried out with approval from Vanderbilt University and Stanford University Offices of Sponsored Research.

Across all participants, 276 individuals completed at least one concurrent neuroimaging and behavioral session. Of these participants, 63 completed just one session, 59 completed two sessions, 97 completed three sessions, and 57 completed all four behavioral and neuroimaging sessions. Because we are interested in examining the within-individual dynamics of white matter, reading, and math development, we limited our analyses to participants who completed three or more sessions, as these dynamic systems cannot be studied with two or fewer observations per participant. Furthermore, there were seven sibling pairs in the dataset. To avoid any potential confounds due to family effects, we randomly excluded one participant from each sibling pair.

After controlling for image quality (see *Neuroimaging Data* for overview on quality control procedures), we had a final sample of 101 participants (52 female participants, and 49 male participants) with the necessary behavioral and neuroimaging data at a minimum of three time points. For our longitudinal analyses, we examined Woodcock-Johnson W scores. The Woodcock-Johnson W score is a mathematical transformation of the Rasch Model on an equal-interval scale (Woodcock & Dahl, 1971; Woodcock, 1999), meaning that they are particularly well-suited for examining within-individual growth in a given metric. At the first time point, participants were, on average, 7.45 years old (SD = 0.34) and displayed an average Woodcock-Johnson W score in Basic Reading of 478.24 (SD = 19.80) and an average Woodcock-Johnson W score in Mathematics of 466.26 (SD = 14.24). The Woodcock-Johnson Basic Reading scores used in this analysis were calculated as the average W score on the word identification and word attack sub-tests and the Woodcock-Johnson Mathematics scores were calculated as the average W score on the calculations and applied problems sub-tests.

### 2.2 Neuroimaging Data

At each time point, MRI data were acquired at Vanderbilt University Institute of Imaging Science on three different 3T Phillips Achieva scanners, each with an 32-channel head coil. During each scan session, diffusion-weighted (DWI) and T1-weighted anatomical images were collected. The DWI scans were acquired using a single-shell echo planar imaging sequence (EPI) with TR/ TE = 8600/66 msec and b =2000sec/mm2 in 60 diffusion directions. Additionally, 6 non-diffusion weighted volumes (b=0) were acquired. The data were acquired in 96 x 94 matrices with an isotropic voxel resolution of 2.5mm^3^. Diffusion weighting was applied along 60 gradient directions evenly distributed across the unit sphere. Acquisition time was 9 min and 6.28 sec per scan.

The T1-weighted images were acquired using a gradient recalled echo protocol (MP-RAGE) with TR/TE =8.9/4.61 msec, flip angle =8°, and an isotropic voxel size of (1 mm)^3^. Each slice was acquired in a 256 x 256 matrix. The total acquisition time was 4min and 24.3sec per scan.

### 2.3 Preprocessing

Preprocessing was performed using QSIPrep 0.16.1, which is based on Nipype 1.8.5 (Gorgolewski et al. (2011); Gorgolewski et al. (2018); RRID:SCR_002502).

### 2.4 Anatomical data preprocessing

The T1-weighted (T1w) image was corrected for intensity non-uniformity (INU) using N4BiasFieldCorrection (Tustison et al. 2010, ANTs 2.4.0), and used as T1w-reference throughout the workflow. The T1w-reference was then skull-stripped using antsBrainExtraction.sh (ANTs 2.4.0), using OASIS as target template. Spatial normalization to the ICBM 152 Nonlinear Asymmetrical template version 2009c (Fonov et al. 2009, RRID:SCR_008796) was performed through nonlinear registration with antsRegistration (ANTs 2.4.0, RRID:SCR_004757, Avants et al. 2008), using brain-extracted versions of both T1w volume and template. Brain tissue segmentation of cerebrospinal fluid (CSF), white-matter (WM) and gray-matter (GM) was performed on the brain-extracted T1w using FAST (FSL 6.0.5.1:57b01774, RRID:SCR_002823, Zhang, Brady, and Smith 2001).

### 2.5 Diffusion data preprocessing

MP-PCA denoising as implemented in MRtrix3’s dwidenoise(Veraart et al. 2016) was applied with a 5-voxel window. After MP-PCA, B1 field inhomogeneity was corrected using dwibiascorrect from MRtrix3 with the N4 algorithm (Tustison et al. 2010). After B1 bias correction, the mean intensity of the DWI series was adjusted so all the mean intensity of the b=0 images matched across each separate DWI scanning sequence.

FSL (version 6.0.5.1:57b01774)’s eddy was used for head motion correction and Eddy current correction (Andersson and Sotiropoulos 2016). Eddy was configured with a q-space smoothing factor of 10, a total of 5 iterations, and 1000 voxels used to estimate hyperparameters. A linear first level model and a linear second level model were used to characterize Eddy current-related spatial distortion. q-space coordinates were forcefully assigned to shells. Field offset was attempted to be separated from subject movement. Shells were aligned post-eddy. Eddy’s outlier replacement was run (Andersson et al. 2016). Data were grouped by slice, only including values from slices determined to contain at least 250 intracerebral voxels. Groups deviating by more than 4 standard deviations from the prediction had their data replaced with imputed values. Final interpolation was performed using the jac method.

Based on the estimated susceptibility distortion, an unwarped b=0 reference was calculated for a more accurate co-registration with the anatomical reference. Several confounding time-series were calculated based on the preprocessed DWI: framewise displacement (FD) using the implementation in Nipype (following the definitions by Power et al. 2014). The head-motion estimates calculated in the correction step were also placed within the corresponding confounds file. Slicewise cross correlation was also calculated. The DWI time-series were resampled to ACPC, generating a preprocessed DWI run in ACPC space with 1.5mm isotropic voxels. Many internal operations of QSIPrep use Nilearn 0.9.2 (Abraham et al. 2014, RRID:SCR_001362) and Dipy (Garyfallidis et al. 2014). For more details of the pipeline, see the section corresponding to workflows in QSIPrep’s documentation (https://qsiprep.readthedocs.io/).

### 2.6 Tractometry with pyAFQ

After preprocessing the diffusion data, we used pyAFQ (Kruper et al., 2021) to perform tractography and calculate tractometry metrics for each participant. Fiber orientation distributions (FOD) were estimated in each voxel using constrained spherical deconvolution implemented in DIPY (Garyfallidis et al., 2014; Tournier et al., 2007). After generating FODs, probabilistic tractography was used to generate streamlines in the white matter. From these streamlines, 22 major white matter tracts were identified using the approach originally outlined by Yeatman et al. (2012). Once identified, each tract was sampled to 100 nodes and fractional anisotropy (FA), mean diffusivity (MD), radial diffusivity (RD), and axial diffusivity (AD) were calculated at each node using a diffusion tensor model (DTI). These tract profile data were then used to examine the relationship between white matter development and academic skills.

### 2.7 Quality Control

To ensure that only high quality neuroimaging data were used in our statistical analyses, we utilized *dmriprep-viewer* to generate quality control scores from the visual reports generated by QSIPrep (Richie-Halford et al., 2022a; Richie-Halford et al., 2022b). This interactive tool allows raters to generate QC ratings based on a various aspects of the preprocessed imaging data, including: the diffusion acquisition time series, motion parameters over the course of the diffusion acquisition, the brain mask and the b=0 to T1w registration, and a 3-d color coded FA map overlaid on the b=0 image. For the present data, two independent raters first completed a training on how to assess the quality of dMRI data using *dmriprep-viewer* before evaluating the same subset of scans (N=174 scans) to ensure reliability across QC ratings. There was 79% agreement in the QC scores given by the two raters in this subset. After this initial round of QC, the raters discussed the scans they did not agree upon to establish a common understanding for how to score images. QC scores were determined using a pass/fail criteria that involved assessing movement within the DWI scan, visually inspecting images for artifacts, examining slice dropout during preprocessing, and evaluating the adequacy of the FA map of each scan acquisition. After this discussion, the remaining 526 scans were divided in half and reviewed by each rater. Overall, 128 scans did not pass quality control, leaving a final sample of 572 quality scans across all four time points to be used for analysis. This quality control procedure left us with a final sample of 101 participants who had completed either 3 or 4 observations with both behavioral and neuroimaging data.

### 2.8 Data Harmonization

The diffusion MRI data used in this study were collected using three different scanners. To account for potential variation between the scanners, we performed ComBat harmonization (Fortin et al., 2017, 2018; Johnson et al., 2007) on the tract profile data, while protecting potential effects of age and reading ability. We did not protect for math ability, as many participants did not complete math assessments at various time points and would therefore be dropped from the harmonization model due to incomplete data. To ensure that these ComBat models did not overfit the data, we fit our models using 5-fold cross-validation, where all of a participant’s data across all time points were included in the same train/test split. This process removes any potential non-biological effects due to scanner differences that might confound subsequent analysis of the tract profile data (Supplemental Figure 2).

### 2.9 GAMM Modeling

We began our analysis by fitting a series of generalized additive mixed models (GAMM; Hastie & Tibshirani, 1987; Lin & Zhang, 1999; Wood, 2011, 2017) to model white matter development along the length of an entire tract. Briefly, GAMMs make use of non-linear smoothing functions to flexibly capture non-linearities in the data, while also allowing for the inclusion of both random effects and parametric fixed effects. These models are especially effective at capturing non-linearities in both space and time, which makes them well-suited for modeling the longitudinal tract profile data generated by pyAFQ. In the present analysis, we fit a series of GAMMs modeling the development of mean diffusivity at each node along the length of a given tract.

In our GAMM examining the link between white matter development and reading, we included initial age, sex, handedness, average reading scores across time points (Reading-Trait), and within-participant centered reading (Reading-State) as parametric terms. We also included smoothing terms on time elapsed since the first observation, position on the tract, and a tract position by reading-state interaction. This model also included random effects of participant, time, and position along the tract to allow for individual differences in overall MD, the rate of MD development, and the variation in MD along the length of the tract. We also incorporated an AR1 model in our GAMM (with ⍴=0.965) to account for the spatial autocorrelations present in tract profile data (Van Rij et al., 2019). The value of ⍴ was determined using the procedure outlined in Van Rij et al. (2019), whereby we fit our model without any autocorrelation structure and then calculated the AFC lag score of the model. We fit this model in both the left and right arcuate to test whether a potential link between growth in reading and white matter development was specific to the left arcuate or a more global phenomenon found throughout the white matter. The GAMM models used in this analysis were specified as follows

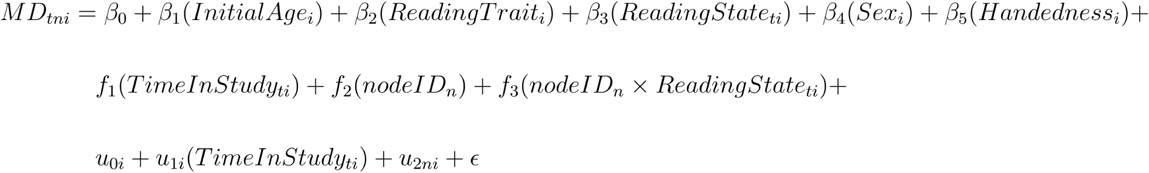

where the *β* coefficients represent the average relationship between the parametric predictors and mean diffusivity at node *n* at time point *t*. Our primary parameters of interest are *β_2_* and *β_3_*. *β_2_* tests the hypothesis that overall reading skill is linked with mean diffusivity whereas *β_3_* tests the hypothesis that within-individual changes in reading are linked to changes in the white matter. The functions *f_i_* represent non-parametric smoothing terms that allow for non-linear relationships between mean diffusivity and the inputs to those functions. *u_0i_* and *u*_1*i*_ represent individual intercepts and growth rates, respectively, and *u_2ni_* represents individual deviations from the prototypical tract profile. Finally, ɛ is an error term representing any variance in mean diffusivity not captured by the model.

To ensure a parsimonious model, we first constructed a GAMM only capturing growth before sequentially adding the reading predictors to the model. At each step, we compared the full and reduced models using a χ^2^ test for nested models and included the predictor only if this test resulted in p-value of less than 0.05. Furthermore, we assessed the adequacy of the number of basis dimensions in each smooth term by examining the significance of the smooth terms of the GAMM predicting the residuals of our proposed model. None of these smooth terms emerged as significant, indicating that the smoothers in our final model are sufficiently “wiggly” and adequately capturing the variance in the non-linear relationships in the data.

### 2.10 Linear Modeling

To examine the longitudinal relationship between the development of the left arcuate and reading development, we fit a linear-mixed effects model predicting mean-centered reading scores, as outlined in Roy et al. (2024). This model included fixed-effects of time point, initial age, within-individual mean-centered mean MD in the left arcuate at each time point (MD-state), and overall mean MD in the left arcuate across all time points (MD-trait). This model also included participant specific random intercepts and slopes on time point to allow for interindividual differences in initial reading scores and growth in reading over time. In addition to this model, we also fit a similar model using a continuous measure of time (time elapsed since the participants initial visit). These models were specified as follows:

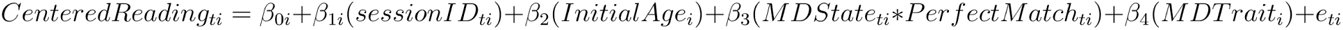

where each participant’s reading score at time *t* is modeled as a function of a participant specific intercept (*β_0i_*), a participant-specific slope (*β*_1*i*_), group effects of initial age (*β_2_*), MD-state (*β_3_*), MD-trait (*β_4_*), and a residual error term (e*_ti_*). The participant-specific parameters, *β_0i_* and *β*_1*i*_ were modeled as:

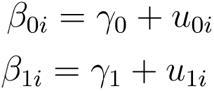

Where γ_0_ and γ_1_ refer to the average initial reading score and average rate of reading development, respectively, and *u_0i_* and *u*_1*i*_ describe the extent to which each individual differs from these averages.

### 2.11 Modeling time series with mlVAR

To model the relationship between the time series of changes in mean diffusivity in the left arcuate and changes in reading scores, we fit a multi-level vector autoregression model (Bringmann et al., 2013; Epskamp, Waldorp, et al., 2018). Briefly, this model leverages the time series data of average mean diffusivity and reading scores to model the temporal dynamics of these two change processes, while accounting for individual differences through random effects structures. From these time series, we can then determine the extent to which reading measures at one time point predict future changes in the white matter and vice versa.

### 2.12 Software and Code Availability

All statistical analyses were conducted using the R programming language (version 4.4.1; R Core Team, 2022). Linear mixed-effects models were fit using the *lme4* package (Bates et al., 2015), GAMM models were fit using the *bam* function present in the *mgcv* package (S. Wood, 2015) with random effect smoothers, and the mlVAR model was fit using the *mlVAR* package (Epskamp, Borsboom, et al., 2018; Epskamp, Waldorp, et al., 2018)

## 3 Results

### 3.1 Within-Individual changes in reading skill track changes in the white matter properties of the left arcuate

We fit a series of generalized additive mixed models (GAMM) to understand how the longitudinal link between reading and white matter development unfolds over the length of the left arcuate. We first visually examined the longitudinal development of four different diffusion metrics calculated by pyAFQ in the left arcuate fasciculus. Supplemental Figure 1 shows how, on average, tract profiles for each diffusion metric develop over time. For the sake of consistency with past studies (Huber et al., 2018), we focus on MD as our main diffusion metric of interest.

Our model predicting MD development in the left arcuate included parametric terms for initial age, mean reading scores across all observations (reading trait), within-individual changes in reading scores (reading-state), sex, and handedness, and smooth terms over the nodes, time, and a node by reading-state interaction. Reading-state and trait were determined using Woodcock-Johnson W scores in Basic Reading (see Methods for overview on Woodcock-Johnson W scores). This model also included a random intercept for each participant and random slopes of node and time, allowing both the tract profile and developmental trajectory of the tract to vary for each participant. Time was operationalized as the amount of time elapsed (in years) from the onset of the experiment. See Methods section for overview of model selection procedure and evaluation of model fit.

The full output of this model is reported in Table 1. The parametric terms of the model suggest that, on average, within-individual gains in reading are coupled to decreases in MD in the left arcuate, while controlling for initial age, sex, handedness, and mean reading score (t(1) = -2.081, p = 0.037; Figure 1A). There was no observed relationship between mean reading scores and MD (t(1) = -0.397, p = 0.691; Figure 1C). The smooth terms of the model revealed a significant non-linear relationship between node (i.e., location on the tract) and MD (F(13.68) = 163.647, p < 0.001), indicating that MD varies over the length of the left arcuate. Additionally, the smooth term on time did not reveal a significant non-linear relationship between time and MD (F(1.884) = 2.607, p = 0.137). However, visual inspection of the smooth estimate revealed that MD tended to decrease over time (Figure 1B), in line with expectations from other developmental studies (Lebel et al., 2012; Moura et al., 2016). There was also no observed smoother interaction between within-individual reading and position along the tract (F(3.468) = 2.049, p = 0.068), suggesting that the observed parametric relationship between within-individual reading and MD is consistent across the length of the tract.

**Figure 1.**
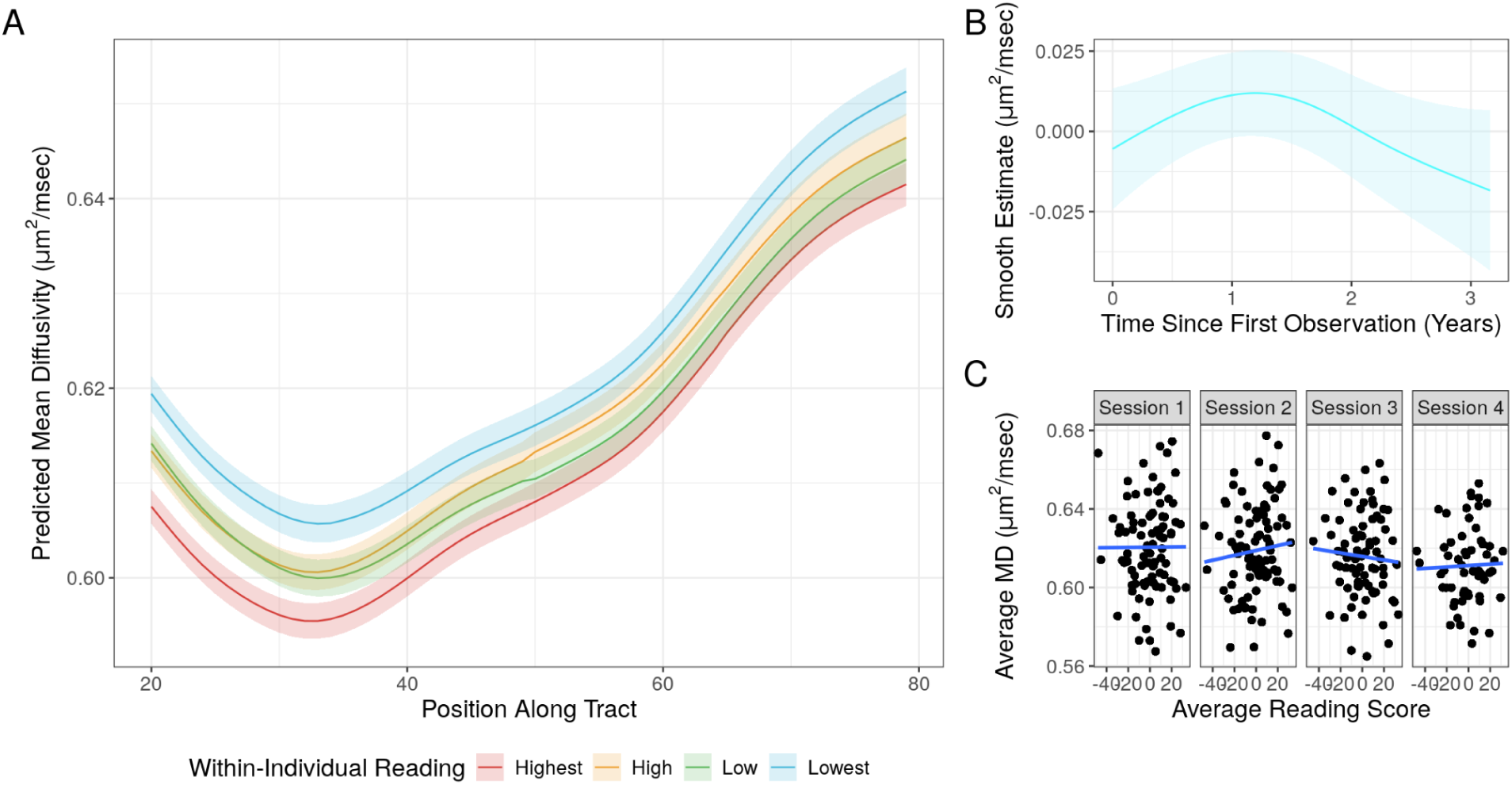
**A.** Average estimated tract profiles for MD in the left arcuate fasciculus generated by the GAMM for four different quartiles of reading score change (reading state). **B.** The estimated smoothing effect of time elapsed since the first study observation on average MD in the left arcuate. This effect was not significant (p = 0.137) **C.** Relationship between overall mean Woodcock-Johnson reading scores and MD in the left arcuate at each time point in the study.

**Table 1:** Summary of parametric coefficients and smooth terms for the final GAMM modeling the development of mean diffusivity (MD) across the length of the left arcuate.

| Component | Term | Estimate | Std. Error | t-value | p-value |
| --- | --- | --- | --- | --- | --- |
| Parametric Coefficients | (Intercept) | 5.974e-04 | 3.511e-05 | 16.876 | <0.001*** |
|  | Initial Age | 2.510e-06 | 4.652e-06 | 0.540 | 0.589 |
|  | Mean Reading | -4.668e-08 | 1.175e-07 | -0.397 | 0.691 |
|  | Centered-Reading | -2.031e-07 | 9.757e-08 | -2.081 | 0.037** |
|  | Sex | 1.098e-05 | 3.986e-06 | 2.755 | 0.006** |
|  | Handedness | 5.345e-08 | 4.374e-08 | 1.192 | 0.233 |
| Component | Term | edf | Ref. df | F-value | p-value |
| Smooth Terms | s(Time) | 1.884 | 2.271 | 2.607 | 0.137 |
|  | s(Node) | 13.682 | 14.794 | 163.647 | <0.001*** |
|  | ti(Node,Centered-Reading) | 3.468 | 5.075 | 2.049 | 0.068 |

Signif. codes: 0 <= '\*\*\*' < 0.001 < '\*\*' < 0.01 < '\*' < 0.05
Adjusted R-squared: 0.809, Deviance explained 0.723
fREML: 4594.2, Scale est: 1.000, N: 19260

To test that the observed relationship between within-individual changes in reading score and changes in the white matter were specific to reading growth and not due to broader maturational changes, we fit the same GAMM to model the development of the right arcuate, a tract not typically associated with reading skill (Vandermosten et al., 2012). This model revealed no significant linear relationship between any of the parametric terms and MD, including within-individual reading and mean reading scores (Supplemental Table 1; all p > 0.05). However, the smooth terms of this model revealed a significant non-linear pattern of MD change in the right arcuate (F(1.75) = 5.172, p = 0.008). Visual inspection of this smoothing term again suggested that, on average, MD decreases as a function of time (Supplementary Figure 3).

Additionally, we fit a model predicting MD development in the left arcuate from mathematics scores to test that the observed relationship between reading and MD development was specific to reading. To do so, we fit the same GAMM using mean math scores and within-individual changes in Woodcock-Johnson W scores in Mathematics, instead of reading scores. The full results are presented in Supplemental Table 2 but briefly, we found no significant relationships between either overall mathematics score or within-individual changes in mathematics scores and MD in the left arcuate (both p > 0.05).

### 3.2 White matter properties and reading skill, but not mathematics, demonstrate between-individual variance in development over a four-year period

We then sought to understand why we observed a longitudinal relationship between the development of the left arcuate and growth in reading, but not math. If there is not significant variance in the growth rates of either white matter maturation or academic skills, then we cannot expect to see a longitudinal, within-individual relationship between these factors. To examine interindividual differences in growth trajectories of academic skills, we fit a series of nested linear-growth models (Grimm et al., 2016) modeling the growth of either Woodcock-Johnson W scores in Basic Reading or Woodcock-Johnson W scores in Mathematics. To identify between-individual differences in the growth trajectories of the MD tract profiles in the left arcuate, we again relied on GAMMs instead of linear mixed-effects models.

In the case of reading, the addition of a term capturing participant-specific developmental trajectories significantly improved the model fit compared to a reduced model that included a random intercept but not a random slope (χ^2^(2) = 19.738, p < 0.001). This finding indicates that there is significant variation in how individuals reading scores change over time (Figure 2A). Interestingly, this was not the case for Woodcock-Johnson W math scores, where the addition of a participant-specific developmental slope did not improve the model fit compared to a reduced model (χ^2^(2) = 0.969, p = 0.616; Figure 2B), suggesting that the participants in this study progressed in mathematics at roughly the same rate.

**Figure 2.**
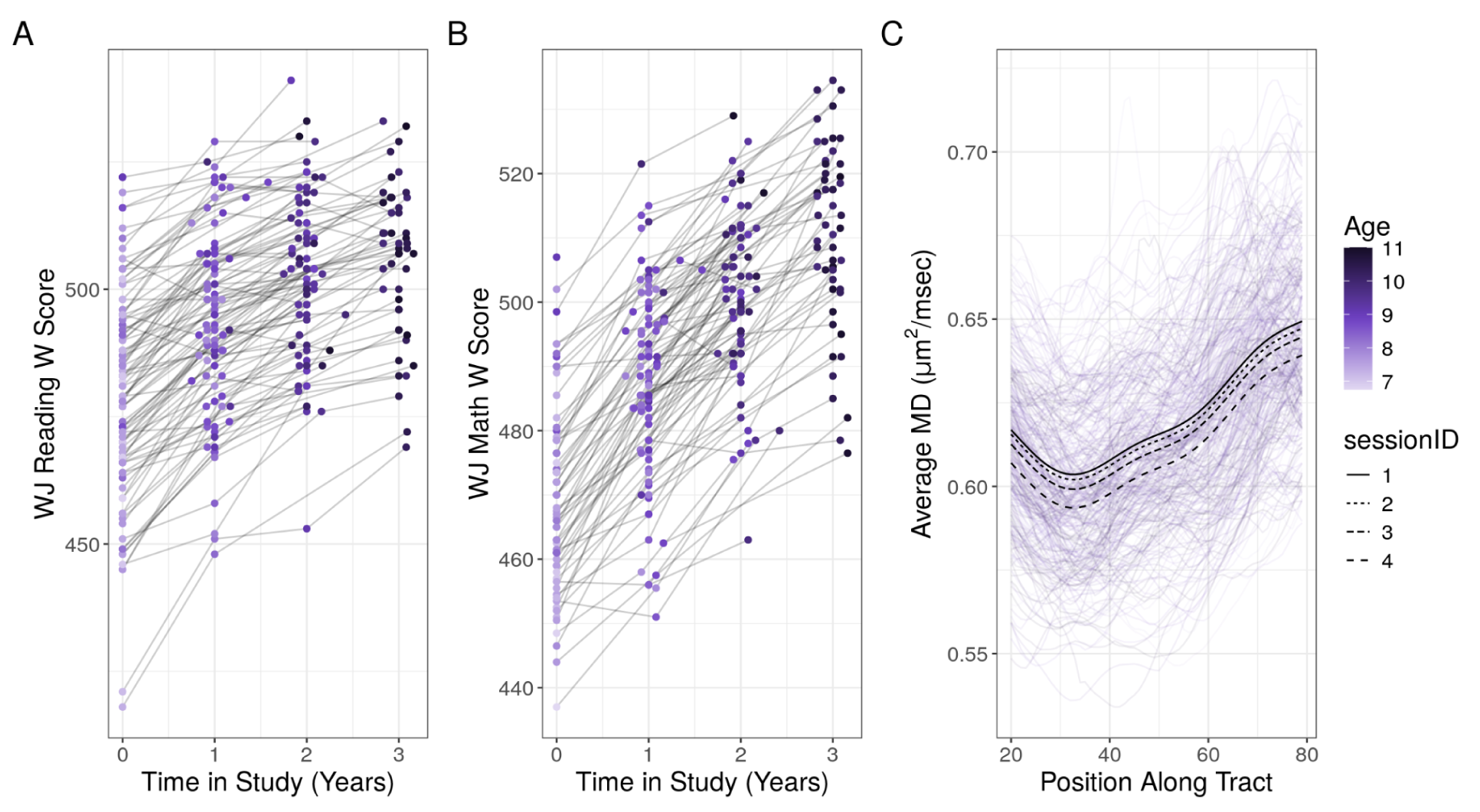
Individual growth trajectories of Woodcock-Johnson composite reading (**A.**) and mathematics (**B.**) W scores. Each line represents an individual participant and the color of each dot represents the participant’s age at a given observation. **C.** Within-individual changes in MD in the left arcuate fasciculus. Each purple profile represents a given participant at a given scan session. The black lines represent the average estimated tract profile at each time point in the study.

To understand between-individual differences in MD development in the left arcuate, we fit a GAMM that included a smooth term over the length of the tract, as well as random effects of participant, position along the tract, and time elapsed since the first observation. The addition of a random slope on time significantly improved the model fit (F(8) = 8.165, p < 0.001), suggesting that there are significant between-individual differences in how MD in the left arcuate changes over time (Figure 2C, Supplemental Figure 2).

### 3.3 Longitudinal changes, not cross-sectional differences, in reading are related to white matter development in the left arcuate fasciculus

After identifying a relationship between the within-individual development of both reading and MD in the left arcuate, we also looked to replicate the findings reported in Meisler and Gabrieli (2022) and Roy et al. (2024) that suggest that static properties (i.e., cross sectional correlations at a given time point) of the left arcuate are not related to reading skill. When we examined the correlation between average MD across the length of the tract and reading skill at each time point, we found no significant relationship between reading scores and MD at any of the four observations (all *r* < 0.12, all p > 0.05; Figure 1C).

We also looked to replicate the finding reported in Roy et al. (2024) suggesting that within-individual changes in reading skill are related to within-individual changes in the mean values of the diffusion properties of the left arcuate. To do so, we calculated the average MD (MD_avg_) of the left arcuate for each participant at each time point and then fit a linear-mixed effects model predicting mean-centered reading using time point, initial age, mean-centered MD_avg_, and MD_avg_ across time (state- and trait-MD_avg_, respectively) as fixed-effects and time point and participant as random effects. This model revealed significant effects of time point (t(102)=-24.614, p<0.001), and initial age (t(255)=-2.033, p = 0.043), indicating that, as expected, reading scores increase over time.

Mean-centered MD_avg_ was also a significant predictor of reading scores (t(316)= -2.003, p = 0.046; Figure 3A), indicating that session-to-session changes in MD predict changes in reading score above and beyond the progression of time. Furthermore, there was no significant relationship between MD_avg_ and mean reading (t(244)=1.750, p = 0.083, Figure 3C). In addition to this model, we also fit a similar linear-mixed effects model using time elapsed since the start of the study as a continuous measure of time, instead of discrete session numbers. Although the coefficients were largely consistent across the two models, the relationship between mean-centered MD_avg_ and mean-centered reading was not statistically significant in the model that included a continuous measure of time (t(314) = -1.851, p = 0.065).

**Figure 3.**
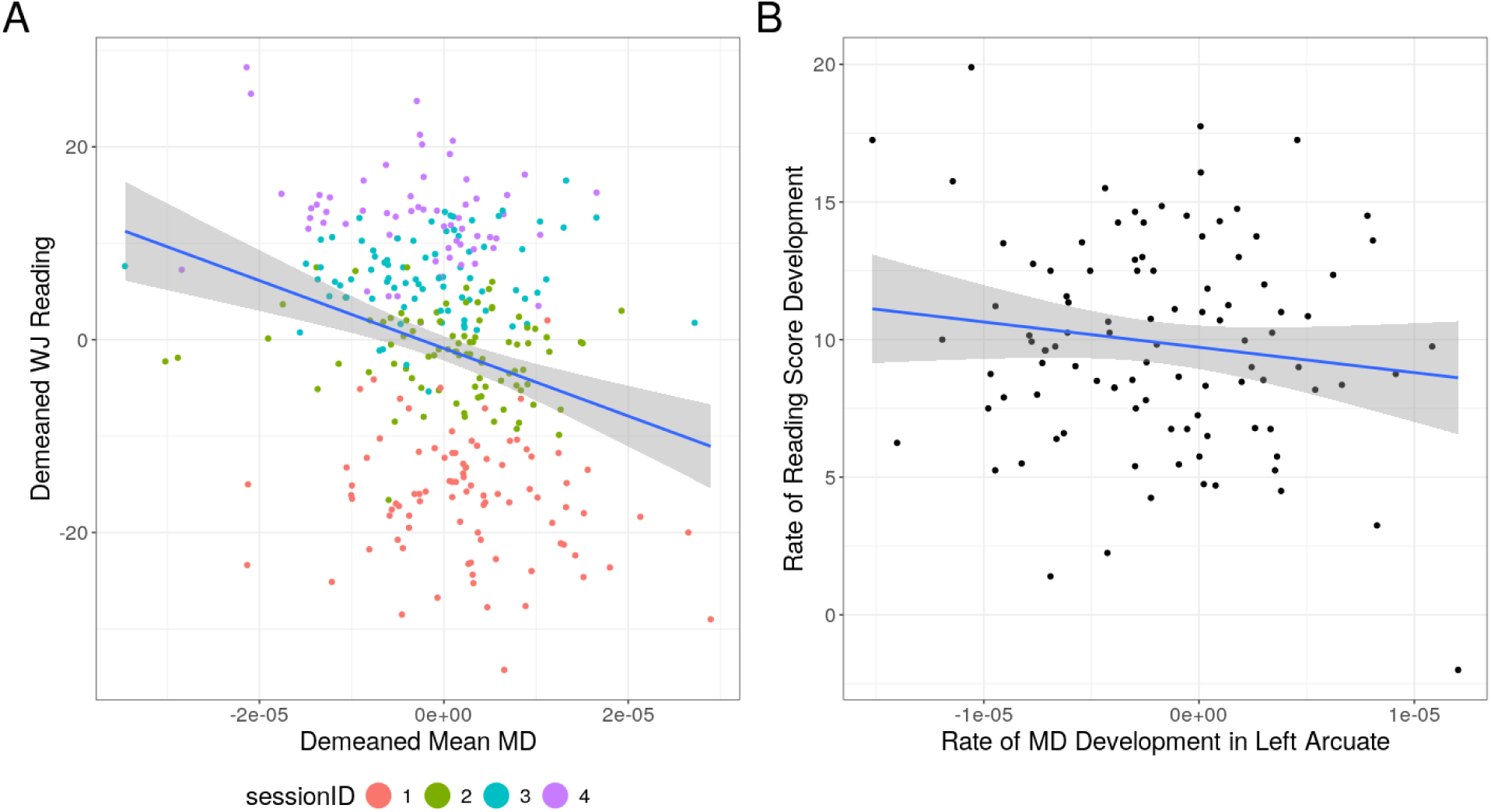
**A.** Relationship between within-individual mean-centered Woodcock-Johnson reading W scores (y-axis) and within-individual mean-centered mean-diffusivity in the left arcuate fasciculus. The color of each dot represents the study session. **B.** Relationship between individual rates of reading development (y-axis) and rate of mean diffusivity (MD) change in the left arcuate (x-axis).

**Figure 4:**
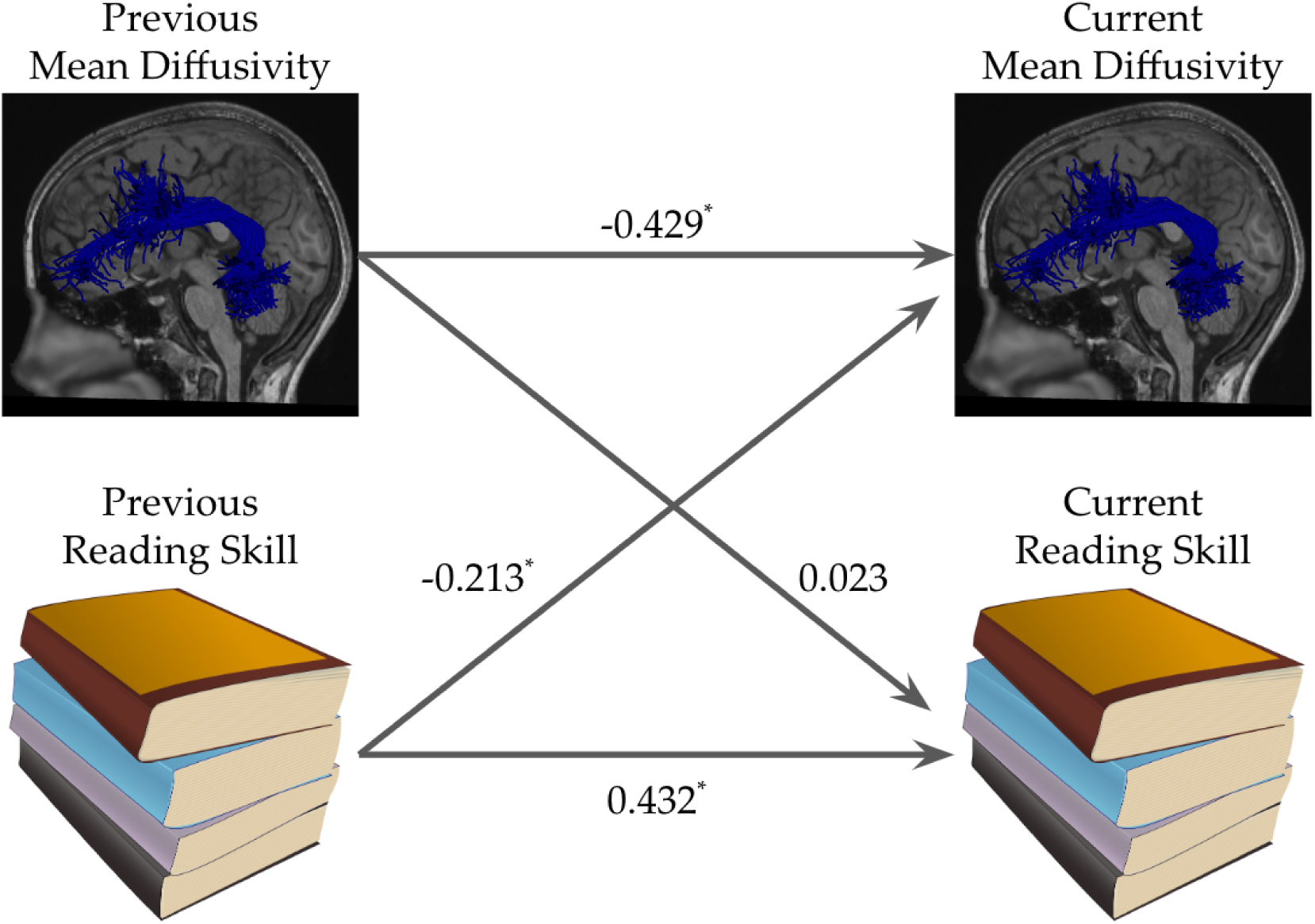
Path diagram illustrating the mlVAR model examining the relationship between the time series of reading development and MD_avg_ in the left arcuate. The values along each path represent the beta-weights estimated by the model. All paths were significant, with the exception of the path of previous MD_avg_ predicting current reading skill.

We then examined the relationship between growth rates in reading and growth rates in MD_avg_ within the left arcuate. To estimate growth rates, we followed the approach outlined in Yeatman et al. (2012) and fit linear models for each participant predicting either reading skill or MD_avg_ as a function of time elapsed from the first observation. We then examined the relationship between the rate of MD_avg_ development and reading development by calculating the correlation between the individual growth rates. We found no significant relationship between the rate of reading growth and MD_avg_ development in the left arcuate (*r* = -0.13, p = 0.19; FIgure 3B), suggesting that, although within-individual changes in reading correspond to within-individual changes in the diffusion properties of the left arcuate, the rate at which these changes occur are not related to one another. Additionally, we examined the variation in MD_avg_ growth rates in relation to average reading scores across all time points to determine whether good versus struggling readers show different patterns of MD_avg_ change in the left arcuate. This analysis revealed no relationship between average reading scores and MD_avg_ development (r = 0.015, p = 0.80 ; Supplemental Figure 4).

### 3.4 Gains in reading precede white matter development across the left arcuate

Given the observed longitudinal relationship between changes in reading scores and changes in MD_avg_ in the left arcuate, we then modeled the relationship between the time series of individual reading development and individual white matter development. To do so, we fit the same multi-level vector auto-regression model (mlVAR; Bringmann et al., 2013; Epskamp et al., 2018) used in Roy et al. (2024). Briefly, these models use the time series of reading scores and white matter development to examine whether growth in one variable predicts future growth in the other, while accounting for between-individual differences through random effects structures. This model revealed that higher reading scores at one time point were associated with lower MD_avg_ at the subsequent time point (t(215) = -2.984, p = 0.003), whereas lower MD_avg_ did not predict future reading scores (t(215) = 0.411, p = 0.681). A bootstrapped difference test (Epskamp, Borsboom, et al., 2018) comparing these two paths revealed that the 95% confidence interval around the difference between these two path weights did not include 0 (95% CI: [0.427, 0.445]), suggesting that these coefficients are significantly different from each other.

## 4. Discussion

In the present study, we leveraged the flexibility of generalized additive mixed-models (GAMM) to understand developmental change in the left arcuate and its relationship to the development of reading skills in a longitudinal sample of 101 children from the Southern United States. Furthermore, we looked to replicate the longitudinal results presented in Roy et al. (2024) in an independent sample and extend them to examine the longitudinal relationship between white matter development and mathematics skill. We employed the same modeling approaches outlined in Roy et al. 2024 to examine the within-individual relationship between white matter development and both reading and math learning.

In the case of reading, we replicated the finding that changes in the diffusion properties of the left arcuate fasciculus, but not the right, correspond to gain in reading over time. At a given time point, we did not observe any cross-sectional relationship between reading skill and white matter properties. However, both the linear and GAMM growth models revealed that within-individual decreases in node-wise MD and MD_avg_ correspond to reading gains over time. Through the use of GAMMs, we were able to extend the original results and account for non-linear developmental trends across the entire length of a given tract. Interestingly, these results revealed that this longitudinal link between MD and reading gains was consistent along the entirety of the left arcuate. Additionally, through the use of mlVAR models, we replicated the finding that gains in reading predict future development of the left arcuate, but not vice versa. Overall, these findings replicate the original findings suggesting a dynamic relationship between reading skill and white matter development that is best viewed through a longitudinal lens.

The observed link between reading growth and white matter development could be due to increased levels of myelination in the reading circuitry, spurred by changes in functional activity. Evidence from animal models has shown that increases in neural activity can increase levels of axonal myelination through an oligodendrocyte-mediated pathway (Fields, 2015; Gibson et al., 2014). In humans, functional MRI studies have demonstrated reduced activity in the ventral occipito-temporal cortex (VOTC) in struggling readers (Brem et al., 2020; Kubota et al., 2019) and that intensive learning interventions can increase both functional activation of VOTC (Heim et al., 2015) and connectivity between this area and other cortical regions (Aboud et al., 2018; Horowitz-Kraus et al., 2019).

Taken together, evidence from animal models and fMRI studies could suggest that intensive reading interventions lead to increased functional activity within the reading circuitry, subsequently driving changes in the white matter properties of these circuits. However, the timescale of these intervention studies do not align well with that reported in the work by Huber et al. (2018), who detected intervention-related changes in the white matter after just 2 weeks. It is likely that different mechanisms of plasticity are at play over different timescales. Future intervention studies combining functional and diffusion neuroimaging data will be necessary to better understand the temporal dynamics of the functional and structural plasticity underlying gains in reading skill.

Despite replicating the longitudinal relationship between session-to-session reading gains and session-to-session white matter development, in the present study we were unable to replicate the original result demonstrating a relationship between the linear rate of reading change and the linear rate of white matter change in the left arcuate. Roy et al. (2024) found that the rate of reading development correlated with the rate of development in the left arcuate and that, when modeled together, the rate of reading development predicts future white matter development. Additionally, Yeatman et al. (2012) found that the rate of development in the left arcuate differed significantly between struggling readers and strong readers. However, these three independent samples come from three distinct geographic areas (the San Francisco Bay Area, the greater San Diego area, and the greater Nashville area) and it could be the case that regional differences in environmental factors, such as educational environment or policy decisions, influence the longitudinal dynamics between reading skill and white matter development (Roy, Van Rinsveld, et al., 2024; Weissman et al., 2023). Future longitudinal work leveraging multi-site consortium datasets will be necessary to better understand this discrepancy.

Furthermore, we extended the longitudinal results presented in Roy, Richie-Halford, et al., 2024 to examine the relationship between the development of the left arcuate and gains in mathematics. However, we did not observe any significant relationships, either cross-sectional or longitudinal, between patterns of white matter development and growth in mathematics. Although we observed intra-individual growth in terms of overall mathematics scores, we did not observe significant interindividual differences in the growth rates of math scores. Additionally, we did not observe a relationship between within-individual growth in math skill and within-individual changes in white matter properties. This finding is not entirely surprising given the lack of variation in growth trajectories in math scores in the current sample.

This finding, however, does not necessarily rule out a longitudinal relationship between mathematics learning and white matter development but rather may reflect a limitation of the present sample. Intervention and quasi-experimental studies have revealed a relationship between educational experiences in mathematics and both functional and structural changes in the brain (Iuculano et al., 2015; Klein et al., 2019). It could be the case that the children in the current sample have all experienced similar educational environments in the domain of mathematics and therefore do not differ significantly in terms of both behavioral gains in mathematics and the underlying white matter changes.

Although generally we have replicated the broad findings linking longitudinal changes in reading skill to changes in the white matter, there are some differences between the present results and those reported in Roy et al. (2024). The primary difference is that, while the previous analysis focused on fractional anisotropy (FA), the present results center on mean diffusivity as the primary white matter metric of interest. Visual inspection of the session by session tract profiles (Supplementary Figure 1) indicated that, in the present sample, FA in the left arcuate does not change across the four time points, whereas mean diffusivity follows the expected developmental trends (Lebel et al., 2019; Lebel & Deoni, 2018). Nevertheless, both FA and MD have been shown to change over the course of development and in response to academic learning experiences (Huber et al., 2018; Klein et al., 2019; Yeatman et al., 2012). The fact that we observed a longitudinal relationship between reading skill and within-participant changes in MD suggest that gains in reading are accompanied by changes in the diffusion properties of the left arcuate. However the biological interpretation is less clear, especially given the surprising lack of developmental change in FA.

In summary, these results largely replicate past findings indicating that changes in the properties of the left arcuate fasciculus are linked longitudinally with gains in reading skill. We extend these findings to a new sample, from a new geographic location, that is weighted towards struggling readers. We also observe that gains in reading precede future changes in the white matter. Together, these results further support the hypothesis that neuroplasticity and academic learning are dynamic processes that are best understood though longitudinal, not cross-sectional, study designs that allow for the study of within-individual change over time.

## Data and Code Availability

The data used for this study are not publicly available but may be made available upon request and completion of a data use agreement. The preprocessed derivatives were generated using the publicly available qsiprep software package (https://qsiprep.readthedocs.io/en/latest/). The pyAFQ outputs were generated using the publicly available pyAFQ software package (https://yeatmanlab.github.io/pyAFQ/). The code used to generate the present analyses and figures can be found at: https://github.com/earoy/longitudinal_read_wm_replication/.

## Author Contributions

**ER**: Conceptualization, Data curation, Formal analysis, Visualization, Writing – original draft, Writing – review & editing. **EMH:** Conceptualization, Data curation, Writing – review & editing **TQN:** Conceptualization, Data collection, Data curation, Writing – review & editing **ARH**: Software, Writing – review & editing. **AR**: Formal analysis, Software, Writing – review & editing. **LEC:** Conceptualization, Data collection, Funding acquisition, Supervision, Writing – review & editing **JDY**: Conceptualization, Formal analysis, Funding acquisition, Supervision, Writing – original draft, Writing – review & editing.

## Supplemental Figures

**Supplemental Figure 1:**
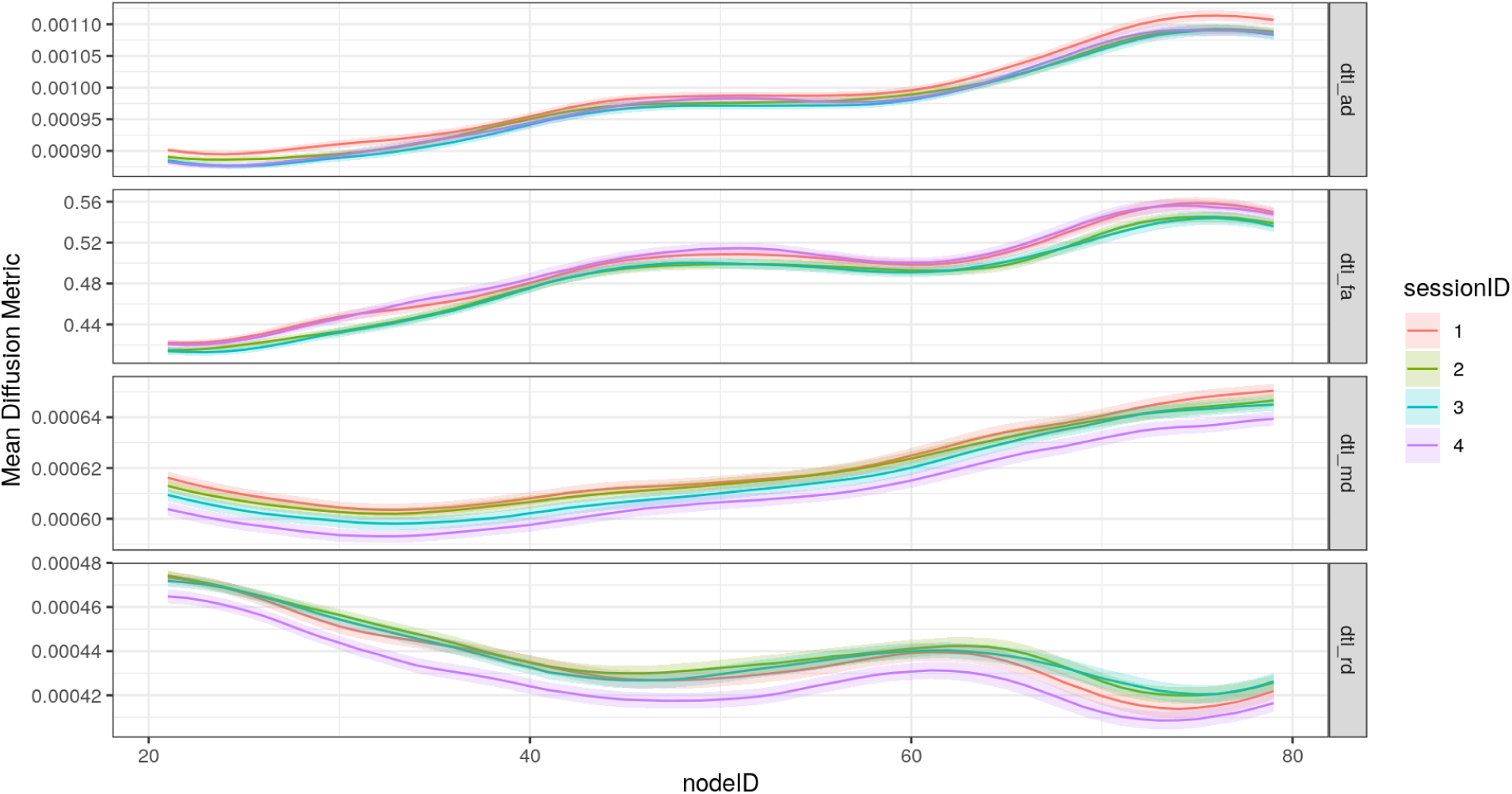
From top, tract profiles in the left arcuate for axial diffusivity, fractional anisotropy, mean diffusivity, and radial diffusivity across the four time points of the study.

**Supplemental Figure 2:**
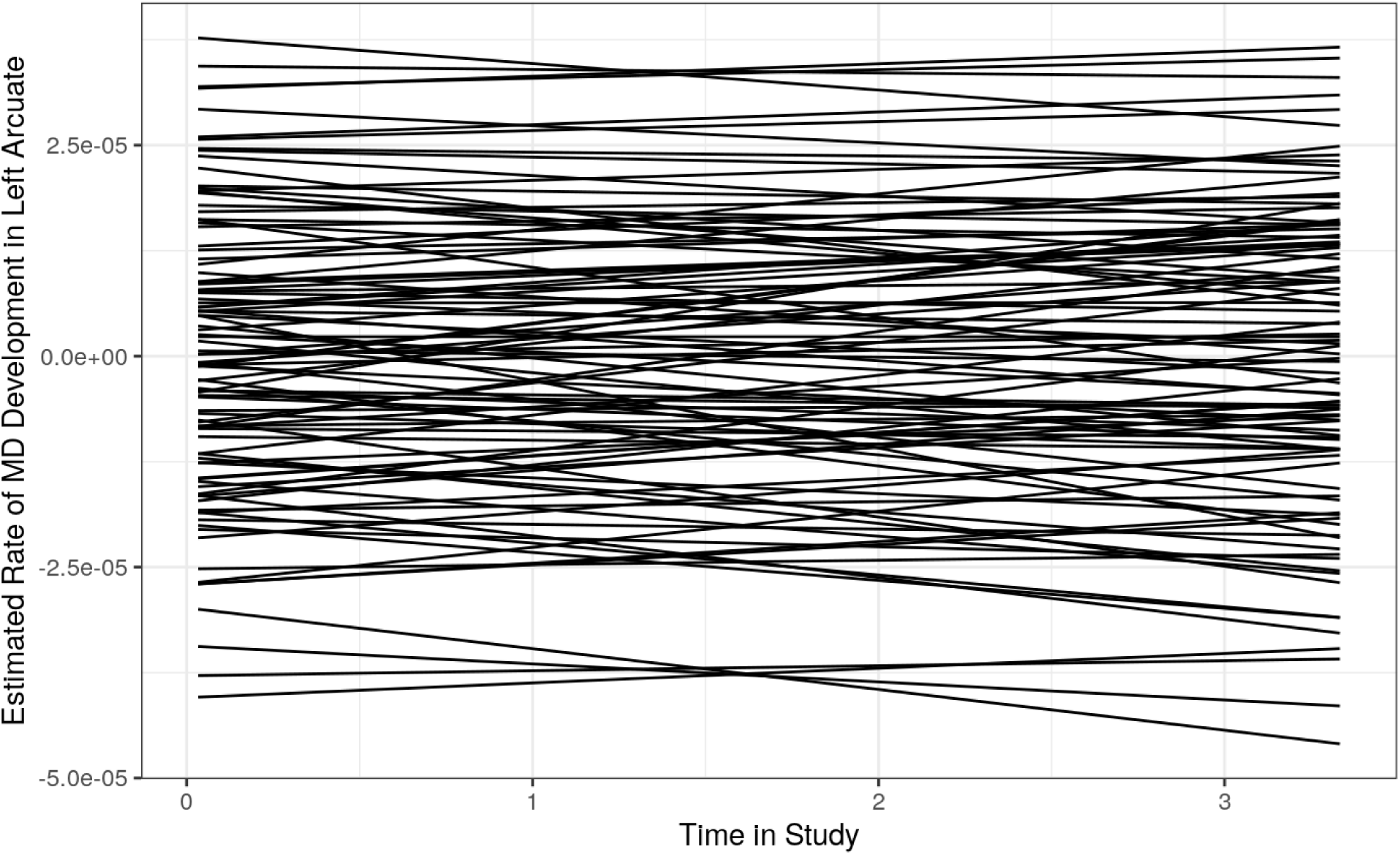
Estimated linear growth rates of MD in the left arcuate. Each line represents the estimated average change in MD over time for each participant.

**Supplemental Figure 3:**
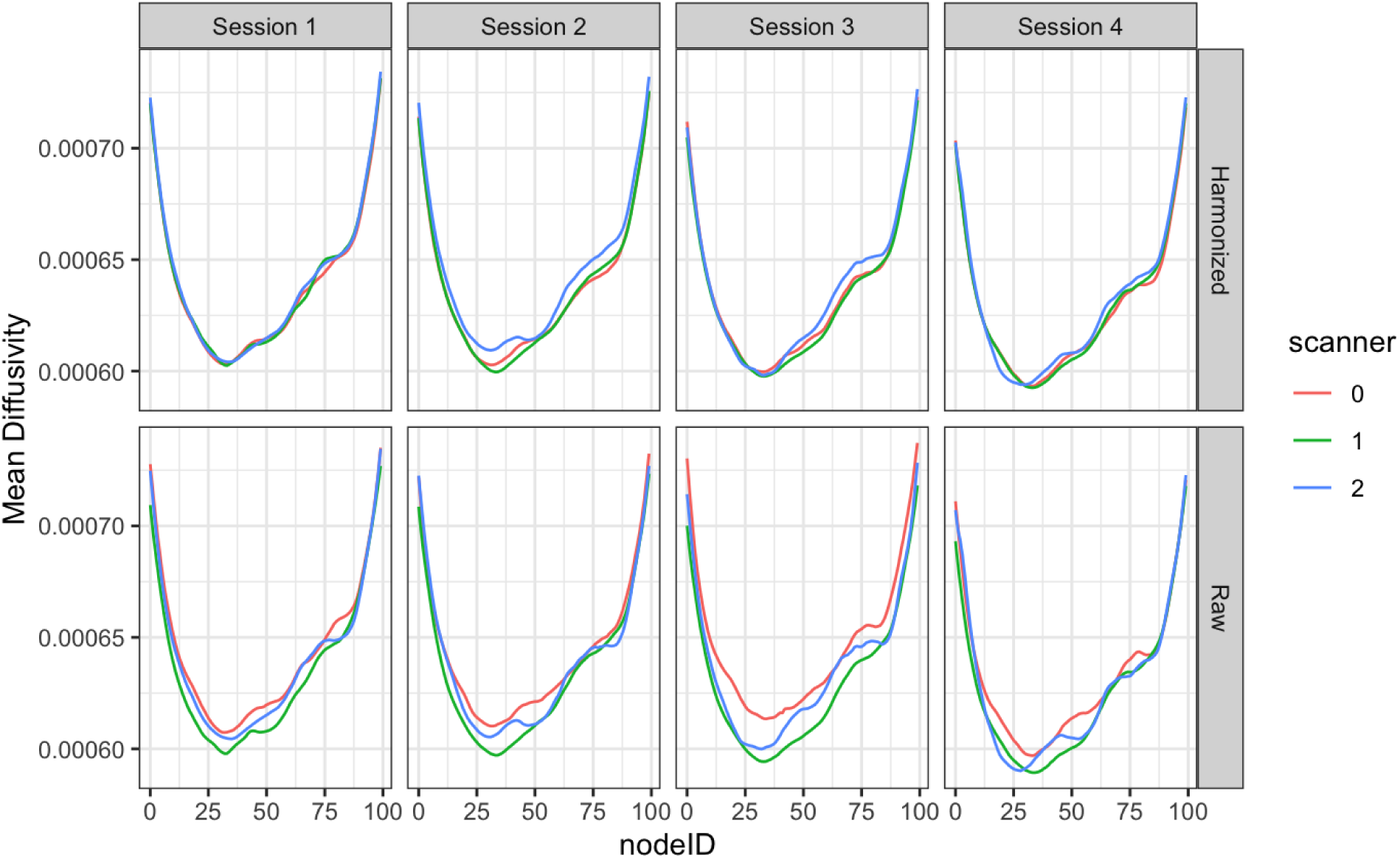
Mean tract profiles illustrating mean diffusivity in the left arcuate using raw tract profile data (bottom row) and ComBat harmonized tract profile data (top row) at each time point. Each color represents a different scanner.

**Supplemental Figure 4:**
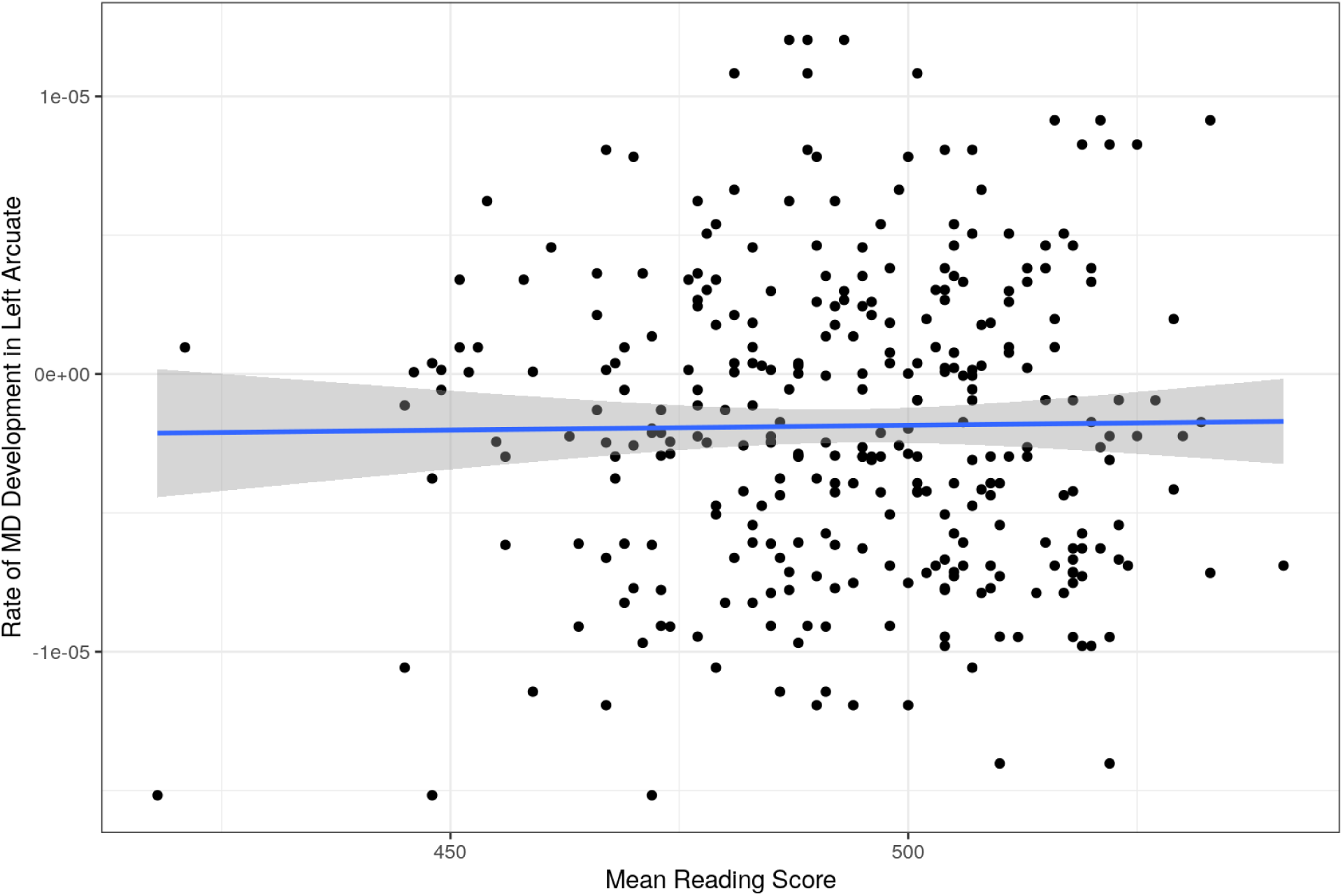
Relationship between average reading scores across all study time points and rates of MD development in the left arcuate for each participant.

**Supplemental Table 1:** Summary of parametric coefficients and smooth terms for the final GAMM modeling the development of mean diffusivity across the length of the right arcuate.

| Component | Term | Estimate | Std. Error | t-value | p-value |
| --- | --- | --- | --- | --- | --- |
| Parametric Coefficients | (Intercept) | 6.14E-04 | 3.63E-05 | 16.902 | <2e-16 *** |
|  | Initial Age | -3.88E-07 | 4.81E-06 | -0.081 | 0.9357 |
|  | Mean Reading | -4.91E-08 | 1.21E-07 | -0.405 | 0.6857 |
|  | Centered-Reading | -7.02E-08 | 9.24E-08 | -0.759 | 0.4478 |
|  | Sex | 9.52E-06 | 4.13E-06 | 2.306 | 0.0211 * |
|  | Handedness | 5.66E-08 | 4.53E-08 | 1.249 | 0.2117 |
| Component | Term | edf | Ref. df | F-value | p-value |
| Smooth Terms | s(Time) | 1.7582 | 2.147 | 5.17E+00 | <0.001*** |
|  | s(Node) | 14.04933 | 14.891 | 2.24E+02 | <0.001*** |
|  | ti(Node,Centered-Reading) | 1.00242 | 1.005 | 5.12E-01 | 0.342 |

Signif. codes: 0 <= '\*\*\*' < 0.001 < '\*\*' < 0.01 < '\*' < 0.05
Adjusted R-squared: 0.821, Deviance explained 0.735
fREML: 5837.8, Scale est: 1.000, N: 19260

**Supplemental Table 2:** Summary of parametric coefficients and smooth terms for the final GAMM modeling the development of mean diffusivity across the length of the left arcuate with math scores.

| Component | Term | Estimate | Std. Error | t-value | p-value |
| --- | --- | --- | --- | --- | --- |
| Parametric Coefficients | (Intercept) | 7.06E-04 | 9.79E-05 | 7.209 | <0.001 *** |
|  | Initial Age | 8.74E-07 | 5.82E-06 | 0.15 | 0.881 |
|  | Mean Math | -2.11E-07 | 2.11E-07 | -1 | 0.317 |
|  | Centered-Math | 5.77E-08 | 1.28E-07 | 0.451 | 0.652 |
|  | Sex | 7.63E-06 | 5.29E-06 | 1.442 | 0.149 |
|  | Handedness | 5.57E-08 | 7.36E-08 | 0.757 | 0.449 |
| Component | Term | edf | Ref. df | F-value | p-value |
| Smooth Terms | s(Time) | 1.00 | 1.00 | 2.71 | 0.107 * |
|  | s(Node) | 12.95 | 14.519 | 1.04E+02 | <0.001 *** |
|  | ti(Node,Centered-Math) | 5.103 | 7.269 | 1.21E+00 | 0.251 |

Signif. codes: 0 <= '\*\*\*' < 0.001 < '\*\*' < 0.01 < '\*' < 0.05
Adjusted R-squared: 0.795, Deviance explained 0.720
fREML: 2631.4, Scale est: 1.000, N: 11340

## References

1. Aboud, K. S., Barquero, L. A., & Cutting, L. E. (2018). Prefrontal mediation of the reading network predicts intervention response in dyslexia. Cortex, 101, 96–106. 10.1016/j.cortex.2018.01.009

2. Bates, D., Mächler, M., Bolker, B., & Walker, S. (2015). Fitting Linear Mixed-Effects Models Using lme4. Journal of Statistical Software, 67(1), 1–48. 10.18637/jss.v067.i01

3. Brem, S., Maurer, U., Kronbichler, M., Schurz, M., Richlan, F., Blau, V., Reithler, J., Van Der Mark, S., Schulz, E., Bucher, K., Moll, K., Landerl, K., Martin, E., Goebel, R., Schulte-Körne, G., Blomert, L., Wimmer, H., & Brandeis, D. (2020). Visual word form processing deficits driven by severity of reading impairments in children with developmental dyslexia. Scientific Reports, 10(1), 18728. 10.1038/s41598-020-75111-8

4. Bringmann, L. F., Vissers, N., Wichers, M., Geschwind, N., Kuppens, P., Peeters, F., Borsboom, D., & Tuerlinckx, F. (2013). A Network Approach to Psychopathology: New Insights into Clinical Longitudinal Data. PLOS ONE, 8(4), e60188. 10.1371/journal.pone.0060188

5. Caffarra, S., Karipidis, I. I., Yablonski, M., & Yeatman, J. D. (2021). Anatomy and physiology of word-selective visual cortex: From visual features to lexical processing. Brain Structure and Function, 226(9), 3051–3065. 10.1007/s00429-021-02384-8

6. Cantlon, J. F., Libertus, M. E., Pinel, P., Dehaene, S., Brannon, E. M., & Pelphrey, K. A. (2009). The Neural Development of an Abstract Concept of Number. Journal of Cognitive Neuroscience, 21(11), 2217–2229. 10.1162/jocn.2008.21159

7. Dehaene, S. (2005). Evolution of human cortical circuits for reading and arithmetic: The “neuronal recycling” hypothesis. From Monkey Brain to Human Brain, 133–157.

8. Dehaene, S., & Cohen, L. (2007). Cultural Recycling of Cortical Maps. Neuron, 56(2), 384–398. 10.1016/j.neuron.2007.10.004

9. Epskamp, S., Borsboom, D., & Fried, E. I. (2018). Estimating psychological networks and their accuracy: A tutorial paper. Behavior Research Methods, 50(1), 195–212. 10.3758/s13428-017-0862-1

10. Epskamp, S., Waldorp, L. J., Mõttus, R., & Borsboom, D. (2018). The Gaussian Graphical Model in Cross-Sectional and Time-Series Data. Multivariate Behavioral Research, 53(4), 453–480. 10.1080/00273171.2018.1454823

11. Fields, R. D. (2015). A new mechanism of nervous system plasticity: Activity-dependent myelination. Nature Reviews Neuroscience, 16(12), Article 12. 10.1038/nrn4023

12. Fortin, J.-P., Cullen, N., Sheline, Y. I., Taylor, W. D., Aselcioglu, I., Cook, P. A., Adams, P., Cooper, C., Fava, M., McGrath, P. J., McInnis, M., Phillips, M. L., Trivedi, M. H., Weissman, M. M., & Shinohara, R. T. (2018). Harmonization of cortical thickness measurements across scanners and sites. NeuroImage, 167, 104–120. 10.1016/j.neuroimage.2017.11.024

13. Fortin, J.-P., Parker, D., Tunç, B., Watanabe, T., Elliott, M. A., Ruparel, K., Roalf, D. R., Satterthwaite, T. D., Gur, R. C., Gur, R. E., Schultz, R. T., Verma, R., & Shinohara, R. T. (2017). Harmonization of multi-site diffusion tensor imaging data. NeuroImage, 161, 149–170. 10.1016/j.neuroimage.2017.08.047

14. Garyfallidis, E., Brett, M., Amirbekian, B., Rokem, A., Van Der Walt, S., Descoteaux, M., Nimmo-Smith, I., & Contributors, D. (2014). Dipy, a library for the analysis of diffusion MRI data. Frontiers in Neuroinformatics, 8, 8.

15. Gibson, E. M., Purger, D., Mount, C. W., Goldstein, A. K., Lin, G. L., Wood, L. S., Inema, I., Miller, S. E., Bieri, G., Zuchero, J. B., Barres, B. A., Woo, P. J., Vogel, H., & Monje, M. (2014). Neuronal Activity Promotes Oligodendrogenesis and Adaptive Myelination in the Mammalian Brain. Science, 344(6183), 1252304–1252304. 10.1126/science.1252304

16. Grimm, K. J., Ram, N., & Estabrook, R. (2016). Growth modeling: Structural equation and multilevel modeling approaches. Guilford Publications.

17. Gullick, M. M., Demir-Lira, Ö. E., & Booth, J. R. (2016). Reading skill–fractional anisotropy relationships in visuospatial tracts diverge depending on socioeconomic status. Developmental Science, 19(4), 673–685.

18. Hastie, T., & Tibshirani, R. (1987). Generalized Additive Models: Some Applications. Journal of the American Statistical Association, 82(398), 371–386. 10.1080/01621459.1987.10478440

19. Heim, S., Pape-Neumann, J., van Ermingen-Marbach, M., Brinkhaus, M., & Grande, M. (2015). Shared vs. Specific brain activation changes in dyslexia after training of phonology, attention, or reading. Brain Structure and Function, 220, 2191–2207.

20. Horowitz-Kraus, T., Hershey, A., Kay, B., & DiFrancesco, M. (2019). Differential effect of reading training on functional connectivity in children with reading difficulties with and without ADHD comorbidity. Journal of Neurolinguistics, 49, 93–108. 10.1016/j.jneuroling.2018.09.002

21. Huber, E., Donnelly, P. M., Rokem, A., & Yeatman, J. D. (2018). Rapid and widespread white matter plasticity during an intensive reading intervention. Nature Communications, 9(1), Article 1. 10.1038/s41467-018-04627-5

22. Iuculano, T., Rosenberg-Lee, M., Richardson, J., Tenison, C., Fuchs, L., Supekar, K., & Menon, V. (2015). Cognitive tutoring induces widespread neuroplasticity and remediates brain function in children with mathematical learning disabilities. Nature Communications, 6(1), 1–10. 10.1038/ncomms9453

23. Johnson, W. E., Li, C., & Rabinovic, A. (2007). Adjusting batch effects in microarray expression data using empirical Bayes methods. Biostatistics, 8(1), 118–127.

24. Jolles, D., Wassermann, D., Chokhani, R., Richardson, J., Tenison, C., Bammer, R., Fuchs, L., Supekar, K., & Menon, V. (2016). Plasticity of left perisylvian white-matter tracts is associated with individual differences in math learning. Brain Structure & Function, 221, 1337–1351. 10.1007/s00429-014-0975-6

25. Klein, E., Willmes, K., Bieck, S. M., Bloechle, J., & Moeller, K. (2019). White matter neuro-plasticity in mental arithmetic: Changes in hippocampal connectivity following arithmetic drill training. Cortex, 114, 115–123. 10.1016/j.cortex.2018.05.017

26. Kruper, J., Yeatman, J. D., Richie-Halford, A., Bloom, D., Grotheer, M., Caffarra, S., Kiar, G., Karipidis, I. I., Roy, E., Chandio, B. Q., Garyfallidis, E., & Rokem, A. (2021). Evaluating the Reliability of Human Brain White Matter Tractometry. Aperture Neuro, 1(1), 10.52294/e6198273-b8e3-4b63-babb-6e6b0da10669. https://doi.org/10.52294/e6198273-b8e3-4b63-babb-6e6b0da10669

27. Kubota, E. C., Joo, S. J., Huber, E., & Yeatman, J. D. (2019). Word selectivity in high-level visual cortex and reading skill. Developmental Cognitive Neuroscience, 36, 100593. 10.1016/j.dcn.2018.09.003

28. Lebel, C., & Deoni, S. (2018). The development of brain white matter microstructure. NeuroImage, 182, 207–218. 10.1016/j.neuroimage.2017.12.097

29. Lebel, C., Gee, M., Camicioli, R., Wieler, M., Martin, W., & Beaulieu, C. (2012). Diffusion tensor imaging of white matter tract evolution over the lifespan. NeuroImage, 60(1), 340–352. 10.1016/j.neuroimage.2011.11.094

30. Lebel, C., Treit, S., & Beaulieu, C. (2019). A review of diffusion MRI of typical white matter development from early childhood to young adulthood. NMR in Biomedicine, 32(4), e3778. 10.1002/nbm.3778

31. Lin, X., & Zhang, D. (1999). Inference in generalized additive mixed models by using smoothing splines. Journal of the Royal Statistical Society Series B: Statistical Methodology, 61(2), 381–400.

32. Matejko, A. A., & Ansari, D. (2015). Drawing connections between white matter and numerical and mathematical cognition: A literature review. Neuroscience & Biobehavioral Reviews, 48, 35–52. 10.1016/j.neubiorev.2014.11.006

33. Matejko, A. A., Price, G. R., Mazzocco, M. M. M., & Ansari, D. (2013). Individual differences in left parietal white matter predict math scores on the Preliminary Scholastic Aptitude Test. NeuroImage, 66, 604–610. 10.1016/j.neuroimage.2012.10.045

34. Meisler, S. L., & Gabrieli, J. D. E. (2021). *A Large-Scale Investigation of White Matter Microstructural Associations with Reading Ability* [Preprint]. Neuroscience. 10.1101/2021.08.26.456137

35. Meisler, S. L., Gabrieli, J. D. E., & Christodoulou, J. A. (2023). *White Matter Microstructural Plasticity Associated with Educational Intervention in Reading Disability* (p. 2023.08.31.553629). bioRxiv. 10.1101/2023.08.31.553629

36. Moreau, D., Wilson, A. J., McKay, N. S., Nihill, K., & Waldie, K. E. (2018). No evidence for systematic white matter correlates of dyslexia and dyscalculia. NeuroImage: Clinical, 18, 356–366. 10.1016/j.nicl.2018.02.004

37. Moura, L. M., Kempton, M., Barker, G., Salum, G., Gadelha, A., Pan, P. M., Hoexter, M., Del Aquilla, M. A. G., Picon, F. A., Anés, M., Otaduy, M. C. G., Amaro, E., Rohde, L. A., McGuire, P., Bressan, R. A., Sato, J. R., & Jackowski, A. P. (2016). Age-effects in white matter using associated diffusion tensor imaging and magnetization transfer ratio during late childhood and early adolescence. Magnetic Resonance Imaging, 34(4), 529–534. 10.1016/j.mri.2015.12.021

38. Muncy, N. M., Kimbler, A., Hedges-Muncy, A. M., McMakin, D. L., & Mattfeld, A. T. (2022). General additive models address statistical issues in diffusion MRI: An example with clinically anxious adolescents. NeuroImage: Clinical, 33, 102937. 10.1016/j.nicl.2022.102937

39. Myers, C. A., Vandermosten, M., Farris, E. A., Hancock, R., Gimenez, P., Black, J. M., Casto, B., Drahos, M., Tumber, M., Hendren, R. L., Hulme, C., & Hoeft, F. (2014). Structural changes in white matter are uniquely related to children’s reading development. Psychological Science, 25(10), 1870–1883. 10.1177/0956797614544511

40. Ozernov-Palchik, O., Norton, E. S., Wang, Y., Beach, S. D., Zuk, J., Wolf, M., Gabrieli, J. D. E., & Gaab, N. (2019). The relationship between socioeconomic status and white matter microstructure in pre-reading children: A longitudinal investigation. Human Brain Mapping, 40(3), 741–754. 10.1002/hbm.24407

41. Polspoel, B., Vandermosten, M., & De Smedt, B. (2019). Relating individual differences in white matter pathways to children’s arithmetic fluency: A spherical deconvolution study. Brain Structure and Function, 224(1), 337–350. 10.1007/s00429-018-1770-6

42. R Core Team. (2022). R: A Language and Environment for Statistical Computing. R Foundation for Statistical Computing. https://www.R-project.org/

43. Ranpura, A., Isaacs, E., Edmonds, C., Rogers, M., Lanigan, J., Singhal, A., Clayden, J., Clark, C., & Butterworth, B. (2013). Developmental trajectories of grey and white matter in dyscalculia. Trends in Neuroscience and Education, 2(2), 56–64. 10.1016/j.tine.2013.06.007

44. Richie-Halford, A., Cieslak, M., Ai, L., Caffarra, S., Covitz, S., Franco, A. R., Karipidis, I. I., Kruper, J., Milham, M., Avelar-Pereira, B., & others. (2022). An analysis-ready and quality controlled resource for pediatric brain white-matter research. Scientific Data, 9(1), 616.

45. Richie-Halford, A. & others. (2022). NiRV: the Neuroimaging Report Viewer. Organization for Human Brain Mapping 2022.

46. Roy, E., Richie-Halford, A., Kruper, J., Narayan, M., Bloom, D., Nedelec, P., Rauschecker, A. M., Sugrue, L. P., Brown, T. T., Jernigan, T. L., McCandliss, B. D., Rokem, A., & Yeatman, J. D. (2024). White matter and literacy: A dynamic system in flux. Developmental Cognitive Neuroscience, 101341. 10.1016/j.dcn.2024.101341

47. Roy, E., Van Rinsveld, A., Nedelec, P., Richie-Halford, A., Rauschecker, A. M., Sugrue, L. P., Rokem, A., McCandliss, B. D., & Yeatman, J. D. (2024). Differences in educational opportunity predict white matter development. Developmental Cognitive Neuroscience, 67, 101386.

48. Rykhlevskaia, E., Uddin, L. Q., Kondos, L., & Menon, V. (2009). Neuroanatomical correlates of developmental dyscalculia: Combined evidence from morphometry and tractography. Frontiers in Human Neuroscience, 3. 10.3389/neuro.09.051.2009

49. Sagi, Y., Tavor, I., Hofstetter, S., Tzur-Moryosef, S., Blumenfeld-Katzir, T., & Assaf, Y. (2012). Learning in the Fast Lane: New Insights into Neuroplasticity. Neuron, 73(6), 1195–1203. 10.1016/j.neuron.2012.01.025

50. Till, C., Deotto, A., Tipu, V., Sled, J. G., Bethune, A., Narayanan, S., Arnold, D. L., & Banwell, B. L. (2011). White matter integrity and math performance in pediatric multiple sclerosis: A diffusion tensor imaging study. NeuroReport, 22(18), 1005–1009. 10.1097/WNR.0b013e32834dc301

51. Tournier, J.-D., Calamante, F., & Connelly, A. (2007). Robust determination of the fibre orientation distribution in diffusion MRI: Non-negativity constrained super-resolved spherical deconvolution. NeuroImage, 35(4), 1459–1472. 10.1016/j.neuroimage.2007.02.016

52. Tsang, J. M., Dougherty, R. F., Deutsch, G. K., Wandell, B. A., & Ben-Shachar, M. (2009). Frontoparietal white matter diffusion properties predict mental arithmetic skills in children. Proceedings of the National Academy of Sciences, 106(52), 22546–22551. 10.1073/pnas.0906094106

53. Turesky, T. K., Sanfilippo, J., Zuk, J., Ahtam, B., Gagoski, B., Lee, A., Garrisi, K., Dunstan, J., Carruthers, C., Vanderauwera, J., Yu, X., & Gaab, N. (2022). Home language and literacy environment and its relationship to socioeconomic status and white matter structure in infancy. Brain Structure and Function, 227(8), 2633–2645. 10.1007/s00429-022-02560-4

54. Van Beek, L., Ghesquière, P., Lagae, L., & De Smedt, B. (2014). Left fronto-parietal white matter correlates with individual differences in children’s ability to solve additions and multiplications: A tractography study. NeuroImage, 90, 117–127. 10.1016/j.neuroimage.2013.12.030

55. Van Der Auwera, S., Vandermosten, M., Wouters, J., Ghesquière, P., & Vanderauwera, J. (2021). A three-time point longitudinal investigation of the arcuate fasciculus throughout reading acquisition in children developing dyslexia. NeuroImage, 237, 118087. 10.1016/j.neuroimage.2021.118087

56. van Eimeren, L., Niogi, S. N., McCandliss, B. D., Holloway, I. D., & Ansari, D. (2008). White matter microstructures underlying mathematical abilities in children: NeuroReport, 19(11), 1117–1121. 10.1097/WNR.0b013e328307f5c1

57. Van Rij, J., Hendriks, P., Van Rijn, H., Baayen, R. H., & Wood, S. N. (2019). Analyzing the Time Course of Pupillometric Data. Trends in Hearing, 23, 2331216519832483. 10.1177/2331216519832483

58. Vanderauwera, J., Wouters, J., Vandermosten, M., & Ghesquière, P. (2017). Early dynamics of white matter deficits in children developing dyslexia. Developmental Cognitive Neuroscience, 27, 69–77. 10.1016/j.dcn.2017.08.003

59. Vandermosten, M., Boets, B., Wouters, J., & Ghesquière, P. (2012). A qualitative and quantitative review of diffusion tensor imaging studies in reading and dyslexia. Neuroscience & Biobehavioral Reviews, 36(6), 1532–1552. 10.1016/j.neubiorev.2012.04.002

60. Wang, Y., Mauer, M. V., Raney, T., Peysakhovich, B., Becker, B. L. C., Sliva, D. D., & Gaab, N. (2017). Development of Tract-Specific White Matter Pathways During Early Reading Development in At-Risk Children and Typical Controls. Cerebral Cortex, 27(4), 2469–2485. 10.1093/cercor/bhw095

61. Weissman, D. G., Hatzenbuehler, M. L., Cikara, M., Barch, D. M., & McLaughlin, K. A. (2023). State-level macro-economic factors moderate the association of low income with brain structure and mental health in U.S. children. Nature Communications, 14(1), Article 1. 10.1038/s41467-023-37778-1

62. Welcome, S. E., & Joanisse, M. F. (2014). Individual differences in white matter anatomy predict dissociable components of reading skill in adults. NeuroImage, 96, 261–275. 10.1016/j.neuroimage.2014.03.069

63. Wierenga, L. M., van den Heuvel, M. P., Oranje, B., Giedd, J. N., Durston, S., Peper, J. S., Brown, T. T., Crone, E. A., & The Pediatric Longitudinal Imaging, Neurocognition, and Genetics Study. (2018). A multisample study of longitudinal changes in brain network architecture in 4–13-year-old children. Human Brain Mapping, 39(1), 157–170. https://doi.org/10.1002/hbm.23833

64. Wood, S. (2015). Package ‘mgcv.’ R Package Version, 1(29), 729.

65. Wood, S. N. (2011). Fast stable restricted maximum likelihood and marginal likelihood estimation of semiparametric generalized linear models. Journal of the Royal Statistical Society Series B: Statistical Methodology, 73(1), 3–36.

66. Wood, S. N. (2017). Generalized additive models: An introduction with R. chapman and hall/CRC.

67. Woodcock, R., & Dahl, M. (1971). A common scale for the measurement of person ability and test item difficulty (AGS Paper No. 10). Minneapolis, MN: NCS Pearson. Inc.

68. Woodcock, R. W. (1999). What Can Rasch-Based Scores Convey About a Person’s Test Performance? 1. In The new rules of measurement (pp. 105–128). Psychology Press.

69. Yeatman, J. D., Dougherty, R. F., Ben-Shachar, M., & Wandell, B. A. (2012). Development of white matter and reading skills. Proceedings of the National Academy of Sciences, 109(44), E3045–E3053. 10.1073/pnas.1206792109

70. Yeatman, J. D., Dougherty, R. F., Myall, N. J., Wandell, B. A., & Feldman, H. M. (2012). Tract Profiles of White Matter Properties: Automating Fiber-Tract Quantification. PLOS ONE, 7(11), e49790. 10.1371/journal.pone.0049790

71. Yeatman, J. D., Dougherty, R. F., Rykhlevskaia, E., Sherbondy, A. J., Deutsch, G. K., Wandell, B. A., & Ben-Shachar, M. (2011). Anatomical Properties of the Arcuate Fasciculus Predict Phonological and Reading Skills in Children. Journal of Cognitive Neuroscience, 23(11), 3304–3317. 10.1162/jocn_a_00061

